# Filtration attenuates cannabis smoke-induced impacts on primary human airway epithelial cells

**DOI:** 10.64898/2026.11.18.767664

**Authors:** Guanghan Shi, Ryan Singer, Nahal Emami Fard, Tracey Campbell, Nicola Henriquez, Imran Satia, James MacKillop, Roma Sehmi, Nadia Milad, Jeremy A. Hirota

**Affiliations:** Firestone Institute for Respiratory Health, Division of Respirology, Department of Medicine, McMaster University & St. Joseph’s Healthcare, Hamilton, ON, Canada; Centre for Microbial Chemical Biology, McMaster University, Hamilton, ON, Canada; Department of Psychiatry and Behavioural Neurosciences, McMaster University.; Peter Boris Centre for Addictions Research, McMaster University & St. Joseph’s Healthcare; Centre for Medicinal Cannabis Research, McMaster University, Hamilton, ON, Canada; School of Biomedical Engineering, McMaster University, Hamilton, ON, Canada; Division of Respiratory Medicine, Department of Medicine, University of British Columbia, Vancouver, BC, Canada; Department of Biology, University of Waterloo, Waterloo, ON, Canada

**Keywords:** air-liquid interface, cellulose acetate, barrier function, cannabinoid capture, IL-33, mucociliary differentiation, harm reduction

## Abstract

**Rationale:** Combustion of cannabis flower and inhalation into the lungs exposes the airway epithelium to toxic particulates and gases. Whether cigarette-style cellulose-acetate filtration, a standard for tobacco cigarettes, attenuates epithelial injury and how it affects cannabinoid capture remain unclear.

**Methods:** Primary human bronchial epithelial cells from four donors were differentiated at air-liquid interface for 30 days and exposed to three cycles of air or CBD- (12.5%) or THC-dominant (20.2%) cannabis smoke generated with or without a cellulose-acetate filter. Barrier function, epithelial denudation, mediator release and epithelial cell protein markers were assessed. Cannabinoids captured from combustion trials of CBD and THC-dominant cannabis were quantified by LC-MS/MS.

**Results:** Repeated exposure to THC or CBD-dominant cannabis strains decreased barrier function, increased epithelial cell denudation, and resulted in greater cell death. Filtration significantly impeded changes in barrier function, epithelial denudation, and cell-death after exposure to both cannabis products. Filtration reduced THC smoke-induced IL-33, Fas, and TRAIL. Soluble CBD and THC concentrations did not differ significantly between filtered and unfiltered combustion. Despite reducing acute epithelial cell injury and mediator release, filtered smoke did not prevent changes in epithelial cell marker expression profiles.

**Conclusion:** Cellulose-acetate filtration attenuated acute cannabis-smoke-induced epithelial injury and selected mediator responses but did not prevent changes in epithelial cell marker expression profiles associated with differentiated cell states. Measured soluble cannabinoid concentrations did not differ significantly between filtered and unfiltered combustion experiments. Cellulose-acetate filters therefore present a promising harm-reduction strategy for cannabis flower combustion, although it should be noted that like tobacco smoke, negative impacts of cannabis combustion are likely to persist even with filtration.

## Introduction

Cannabis is the most widely used controlled drug worldwide, with an estimated 256 million people using it in 2024 [1]. Canada legalized non-medical cannabis in 2018, and 26% of people aged 16 years or older reported past-year non-medical use in 2024 [2]. Although product preferences have diversified since legalization, smoking remained the most common method of consumption (69%) and dried flower/leaf the most frequently used product (63%) [2]. In a longitudinal Ontario cohort, use of dried flower/leaf declined over the first 5 years after legalization but remained common among active consumers [3]. Combustion therefore continues to represent a major route of cannabis exposure and an important respiratory-health concern.

Legal cannabis products sold in Canada are subject to federal production, contaminant-testing and cannabinoid-labelling requirements [4]. Comparable pre-sale testing frameworks operate in established US legal markets: California requires batch testing for cannabinoid and terpene content and for pesticides, heavy metals, microbial impurities and other contaminants [5], while Colorado regulates production-batch potency and contaminant testing through licensed facilities [6]. These systems improve product characterization but do not establish any measure of respiratory safety after combustion.

Commercial cannabis flower is commonly labelled as indica, sativa or hybrid and is further differentiated by cannabinoid composition, including THC-dominant, CBD-dominant and mixed THC/CBD products. However, indica-sativa labels show limited genome-wide differentiation and do not reliably capture overall phytochemical similarity [7]. Cannabinoid-dominant chemotypes therefore provide a more informative basis for controlled experimental comparisons. The diversity of products available in legal markets also creates a need to define the composition of plant material used in exposure studies.

Regardless of cannabinoid composition, cannabis combustion rapidly delivers cannabinoids but also generates particulate matter, carbon monoxide, aldehydes, polycyclic aromatic hydrocarbons and other thermal degradation products [8]. Regular cannabis smoking is associated with cough, sputum production, wheeze and chronic bronchitic symptoms [9,10]. In the recent CANUCK study, cannabis-smoking participants had worse respiratory health status, reflected by higher St George’s Respiratory Questionnaire (SGRQ) total scores, together with lower pre-bronchodilator FEV1/FVC and FEF25-75, and altered airway epithelial immune signatures and MUC5AC expression compared with never-smokers [11]. *In vitro* studies using primary human airway epithelial cells similarly demonstrate pro-inflammatory and barrier responses to cannabis smoke exposure [12]. These observations support evaluation of interventions that reduce epithelial exposure to harmful combustion products.

The airway epithelium is the first tissue contacted by inhaled cannabis smoke. Tight junctions, adherens junctions and desmosomes maintain barrier integrity, while basal, club, goblet and ciliated cells coordinate repair, innate immune signalling and mucociliary clearance [13–15]. In a tobacco-smoke model, cigarette-smoke extract exposure during air-liquid interface (ALI) differentiation impaired barrier formation, reduced acetylated-alpha-tubulin-positive ciliated cells, increased club and goblet cells, and increased expression of the basal-cell marker KRT14 [15]. These findings demonstrate that repeated smoke exposure, using tobacco as a model, can disrupt both barrier function and airway epithelial cell phenotype and differentiation state. This work provides a rationale for determining whether cannabis smoke causes similar injury and airway epithelial cell phenotypic remodelling.

ALI cultures differentiate primary airway epithelial cells into a pseudostratified mucociliary epithelium that more closely recapitulates *in vivo* airway architecture and cellular diversity than immortalized or submerged monolayers [14,16]. The ALI format supports apical smoke exposure, serial barrier function measurements, direct imaging, basolateral mediator sampling and epithelial phenotyping. Our group has developed open-source Transwell exposure systems [17,18] and previously demonstrated dose- and cannabis-product-dependent effects on epithelial barrier function [12,18]. However, these studies did not explore mechanisms to reduce the harms of cannabis combustion on airway epithelial cell biology, that may include physical filtering of mainstream cannabis smoke as is done for tobacco cigarettes.

Conventional cellulose-acetate cigarette filters are not standard components of cannabis joints. Such filters became ubiquitous in manufactured cigarettes as a means of reducing machine-measured smoke constituents, but their incomplete protection and contribution to misleading perceptions of safety are well documented [19]. They retain particulate and condensable material, whereas removal of gas-phase constituents varies with filter design and smoking conditions [20,21]. Whether a conventional cellulose-acetate cigarette filter attenuates cannabis-smoke-induced airway epithelial injury without substantially reducing cannabinoid capture is unknown. We therefore compared repeated filtered and unfiltered CBD- and THC-dominant cannabis-smoke exposures in donor-derived ALI cultures and quantified major cannabinoids in experimental samples. We hypothesized that filtration would attenuate cannabis smoke induced barrier disruption, epithelial injury, cell death, mediator release and epithelial phenotypic changes while retaining measurable cannabinoid delivery performance.

## Methods

### Primary cells and ALI culture

Primary human bronchial epithelial cells were obtained from seven de-identified participants enrolled with informed consent under Hamilton Integrated Research Ethics Board protocol 17668. Cells at passages 2-4 were seeded at 100,000 cells per insert on 24-well Transwell inserts (0.4-µm polyester membranes; 0.33 cm²; Corning, cat. no. 3470, Corning, NY, USA) coated with bovine plasma fibronectin (Sigma-Aldrich, cat. no. F4759-2MG, St Louis, MO, USA) and differentiated at ALI for 30 days in PneumaCult-ALI Medium (STEMCELL Technologies, cat. no. 05001, Vancouver, BC, Canada) as previously described [17,18]. One insert per donor and condition was used for Trans-Epithelial Electrical Resistance (TEER), a measurement of barrier function. The same insert per donor condition was paired for imaging. Lactate dehydrogenase (LDH) release and immune mediator profiling were assessed in basolateral cell-culture medium. For flow cytometry, cells from four parallel inserts from the same donor and condition were pooled to generate sufficient cells to represent a single donor sample.

Donor cultures were assessed after 30 days of ALI differentiation, designated ALI day 0 immediately before the first exposure. Quality control required complete epithelial coverage of the insert surface and a mean raw, area-uncorrected resistance >1000 Ω. Four donor cultures (Donor-02, Donor-06, Donor-22 and Donor-29) met these criteria and were included in smoke-exposure experiments; Donor-16, Donor-27 and Donor-28 were excluded before smoke exposure because they did not meet ALI quality-control criteria. Donor characteristics and ALI quality-control outcomes are shown in Table 1.

**Table 1:** Airway epithelial-cell donor characteristics and ALI quality-control outcomes.

| Donor code | Age (years) | Sex | Race/ethnicity | Mean raw resistance at ALI day 0 (Ω) | ALI QC |
|---|---|---|---|---|---|
| Donor-02 | 51 | Male | Caucasian; not Hispanic/Latino | 1,035 | Passed |
| Donor-06 | 62 | Male | Caucasian; not Hispanic/Latino | 1,480 | Passed |
| Donor-16 | 52 | Male | Caucasian; not Hispanic/Latino | 821 | Failed |
| Donor-22 | 31 | Male | Caucasian; not Hispanic/Latino | 1,802 | Passed |
| Donor-27 | 19 | Male | Asian; not Hispanic/Latino | 679 | Failed |
| Donor-28 | 20 | Male | Asian; not Hispanic/Latino | 954 | Failed |
| Donor-29 | 24 | Male | Asian; not Hispanic/Latino | 1,575 | Passed |

Values are mean raw, area-uncorrected resistance measurements obtained at ALI day 0, immediately before the first exposure. ALI QC required complete epithelial coverage and a mean raw resistance >1000 Ω.

### Cannabis Products and Exposure Protocol

Cannabis smoke exposure was generated using filtered or unfiltered smoke from a CBD and a THC dominant strain. The CBD-dominant strain with trade-name, Painted Lady, (Growtown Cannabis, Kemptville, Ontario) was provided with a batch Certificate of Analysis that quantified total CBD as 10.55% w/w and total THC 0.27% w/w. The THC-dominant strain with trade-name, Sweet Venom Lite (Dykstra Greenhouses, St. Catherines, Ontario) was provided with a batch Certificate of Analysis that quantified total THC as 20.24% w/w and total CBD <0.10% w/w) cannabis (**Table 2**). Both Certificate of Analyses were generated by independent third-party analytical labs as required by legislation governing cannabis production and commercial sale in Canada.

**Table 2:** Cannabinoid composition of the cannabis products used for smoke exposure.

| **Cannabinoid (mg·g<sup>-1</sup>)** | **CBD-dominant strain** | **THC-dominant strain** |
|---|---|---|
| Δ9-THC | ND | 20.2 |
| Δ9-THCA | 3.1 | 207.7 |
| CBD | 12.5 | <0.5 |
| CBDA | 106.0 | <1.0 |
| CBDV | ND | <0.5 |
| CBGA | ND | 2.0 |
| CBG | ND | 1.8 |
| THCV | ND | <0.5 |
| CBN | ND | <0.5 |
| CBC | 4.2 | <1.0 |

Values are batch Certificate of Analysis results provided by an independent third party analytical lab with the accompanied product purchase from a Licensed Provider in the Province of Ontario. ND, not detected below the laboratory-specific limit of detection; “<” denotes a result below the reporting threshold.

All cannabis joints were hand-filled (0.75 g) into identical Premier Supermatic regular king-size filtered cigarette tubes (84 mm; Republic Brands, Glenview, IL, USA) (**Figure 1**). For the non-filter condition, the factory filter from the brown/orange terminal region of the tube (**Figure 1A/1B**) was removed and replaced with a cardboard mouthpiece prepared from RAW Original Tips (HBI Canada, Vancouver, BC, Canada) (**Figure 1B**). For the filtered condition, the cardboard mouthpiece was present with the addition of a cellulose-acetate filter isolated from a 2R4F reference cigarette (Center for Tobacco Reference Products, University of Kentucky, Lexington, KY, USA) (approximately 27×7.9 mm) (**Figure 1A**). Filtered and unfiltered joints underwent complete combustion down to the cardboard mouthpiece.

**Figure 1.**
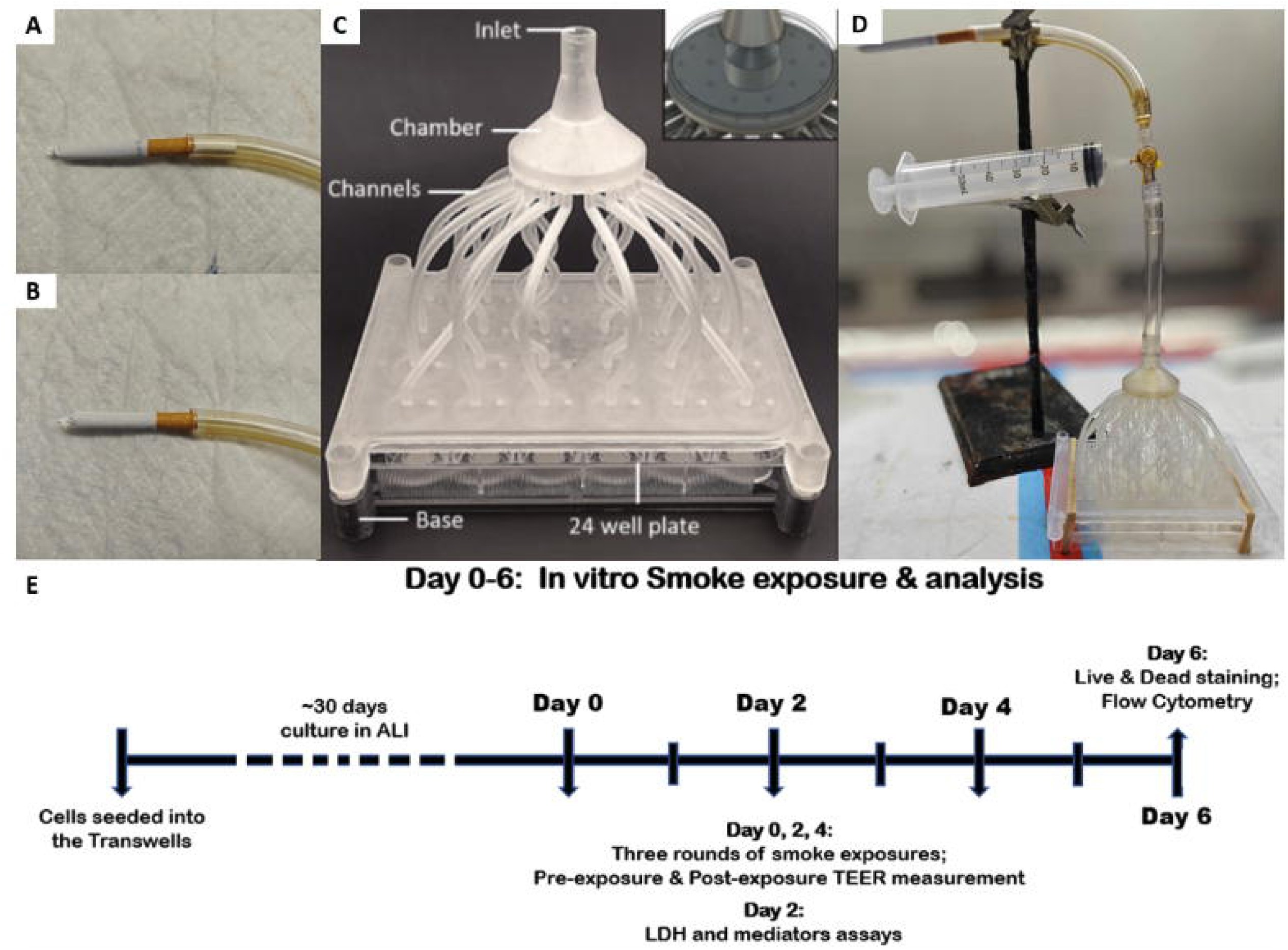
Experimental smoke-exposure system and exposure timeline. (A) Cannabis joint with a cellulose-acetate filter. (B) Joint without a filter. (C) Twenty-four-outlet manifold for ALI cultures [18]. (D) Complete exposure apparatus. (E) Primary human bronchial epithelial cells were differentiated at ALI for approximately 30 days, exposed on days 0, 2 and 4, and assessed on day 6; TEER was measured before and after each exposure.

Mainstream smoke was drawn manually into a sterile 50-mL BD Luer-Lok syringe (BD, cat. no. 309653, Franklin Lakes, NJ, USA) over approximately 10 s and delivered through 40 cm of Tygon S3 E-3603 tubing (Saint-Gobain Performance Plastics, Akron, OH, USA) to a 24-outlet manifold positioned approximately 0.4 cm above the apical surface [18] (**Figure 1C-D**). Each exposure comprised 10 puffs at approximately 20-s intervals over approximately 280 s without air dilution. Air controls followed the same pathway and schedule. Cultures were exposed on days 0, 2 and 4 and assessed on day 6 (**Figure 1E**).

### Airway Epithelial Cell Culture Barrier Resistance and Cell Imaging

TEER was measured immediately before and after exposures on days 0, 2 and 4 and at the day-6 endpoint using a Millicell ERS-2 voltohmmeter (MilliporeSigma, cat. no. MERS00002, Burlington, MA, USA). Blank-insert resistance was subtracted before multiplication by the membrane area, and TEER was expressed as Ω·cm². Values were also normalized to the first pre-exposure measurement. Bright-field images were acquired at the same time points with an EVOS M7000 Imaging System (Invitrogen, cat. no. AMF7000, Carlsbad, CA, USA).

Denuded areas were manually outlined in ImageJ bundled with 64-bit Java 8 (National Institutes of Health, Bethesda, MD, USA) and expressed as a percentage of the Transwell membrane area. After day-6 bright-field imaging, apical surfaces were gently washed with PBS before endpoint processing. At day 6, cultures were stained using the LIVE/DEAD Cell Imaging Kit (488/570; Invitrogen, cat. no. R37601, Carlsbad, CA, USA); ethidium-positive area was quantified using Otsu thresholding and expressed as a percentage of the Transwell membrane area.

### LDH Release and Immune Mediator Profiling

Basolateral medium was collected 48 h after the first exposure, immediately before the second exposure. LDH activity was measured using the CyQUANT LDH Cytotoxicity Assay (Invitrogen, cat. no. C20300, Carlsbad, CA, USA) and expressed as a percentage of maximum release [18]. Immune mediators were measured in undiluted samples using the Human Cytokine/Chemokine Panel B 48-Plex Discovery Assay (Eve Technologies, Calgary, AB, Canada) on a Luminex 200 platform (Luminex Corporation, Austin, TX, USA). The five pre-specified mediators that were primary outcomes presented in the main text were epithelial alarmins (IL-33 and TSLP) and death-associated mediators (soluble Fas, TRAIL and HMGB1). The multi-plex approach was used to maximize the number of mediators that could be measured from the small volume of basal cell culture media collected during experiments.

### Cannabis Smoke Profiling for Cannabinoids

Smoke from each independently combusted 0.75-g CBD- or THC-dominant joint, generated with or without a filter, was bubbled through 4 mL high-glucose DMEM (Gibco, cat. no. 11965-118, Thermo Fisher Scientific, Waltham, MA, USA) using the same puff regimen as the airway epithelial-cell exposures. The capture medium was stored at −20°C until analysis. A 100-μL aliquot was extracted with 400 μL cold LC–MS-grade acetonitrile containing 1% formic acid and matched deuterated internal standards at 50 ng/mL; following protein precipitation, 100 μL of clarified extract was diluted 1:1 with LC–MS-grade water, yielding a final internal-standard concentration of 20 ng/mL. Soluble CBD and Δ9-THC were analysed by positive-ion electrospray multiple-reaction monitoring using an Agilent 1290 Infinity II HPLC coupled to an Agilent 6495C iFunnel triple-quadrupole mass spectrometer and an Agilent RRHD Eclipse Plus C18 column (100×2.1 mm, 1.8 μm).

Quantification used matched d3-labelled internal standards (Sigma-Aldrich, St Louis, MO, USA) and eight-point, 1–200 ng/mL unweighted linear calibration curves. Matrix QC samples prepared at 100, 200 and 1000 ng/mL before extraction underwent the same 10-fold dilution as study samples, yielding final extract concentrations of 10, 20 and 100 ng/mL within the calibration range. Calibration curves required R²>0.99 and back-calculated concentrations within ±30% of nominal; acceptable QC recovery was 80–120% with SD<20%. Study samples and QC samples were injected in duplicate, with a required percentage difference ≤10%; the mean was reported. MDL/LOQ values were 7.5/23 ng/mL for CBD and 9.7/29 ng/mL for THC.

### Flow cytometry

Flow cytometry included air control, filtered CBD smoke and filtered THC smoke experimental conditions; unfiltered conditions were not analysed because severe injury yielded insufficient cells for reliable phenotyping. On day 6, cells from four inserts per donor and condition were pooled to generate sufficient cells to represent a single donor sample. Viability and intracellular acetylated alpha-tubulin (Cell Signaling Technology, cat. no. 18276S, Danvers, MA, USA), MUC5AC (Novus Biologicals, cat. no. NBP2-32732AF647, Centennial, CO, USA), cytokeratin 5 (Abcam, cat. no. ab224984, Cambridge, UK) and SCGB1A1 (Santa Cruz Biotechnology, cat. no. sc-365992 FITC, Dallas, TX, USA) were assessed simultaneously in a single multi-colour cytometry panel. Approximately 200,000–600,000 ungated events were acquired per donor-condition sample on a Cytek Aurora spectral flow cytometer (Cytek Biosciences, Fremont, CA, USA). Unstained, single-stained and antibody-matched isotype controls were included. Using a sequential gating strategy, events were gated on the main cell population, SSC singlets and FSC singlets, followed by FVS700-negative live cells. Marker gates were informed by antibody-matched isotype controls and the observed donor- and condition-dependent background fluorescence. Gates were set at or above the isotype-derived 98th-percentile threshold. If a clear bimodal distribution was present, the interpeak valley was used and the same marker gate was applied across conditions within the donor. Data were analysed in FlowJo v10.8. Events above the final gate were classified as marker^high^, and marker^high^ fractions were calculated among viable singlets and presented as percentages. Representative sequential gating and isotype-control fluorescence distributions are shown in Supplementary Figures S1 and S2.

### Statistical analysis

Analyses were performed in GraphPad Prism 10.5.0. Day-6 TEER was compared between filtered and unfiltered smoke separately within the CBD- and THC-dominant products using two prespecified two-sided paired t-tests. P values were Bonferroni-adjusted across the two comparisons. Denudation, ethidium-positive area and LDH release were analysed across the five exposure conditions using one-way repeated-measures ANOVA without assuming sphericity, with Geisser–Greenhouse correction. Four prespecified Bonferroni-adjusted comparisons were performed for each outcome: air versus unfiltered CBD, air versus unfiltered THC, unfiltered versus filtered CBD, and unfiltered versus filtered THC. All mediator concentrations were positive and were analysed using a lognormal repeated-measures model, consistent with the distributional characteristics of the data and established approaches for multiplex immunoassay measurements [22].

Flow-cytometric outcomes were analysed separately for each marker across air, filtered CBD and filtered THC conditions using one-way repeated-measures ANOVA without assuming sphericity, with Geisser–Greenhouse correction. Two Bonferroni-adjusted comparisons were performed for each marker: filtered CBD versus air and filtered THC versus air. Soluble CBD and THC concentrations were compared between filtered and unfiltered conditions using two-sided unpaired Welch’s t-tests.

Data are presented as individual values with mean±SD. All tests were two-sided. Bonferroni-adjusted p<0.05 was considered statistically significant; significant comparisons were reported as p<0.05.

Additional experimental procedures and analytical details are provided in the online Supplementary Methods.

## Results

### Filtration attenuated cannabis-smoke-induced epithelial barrier injury

Repeated unfiltered CBD- and THC-dominant smoke caused rapid reductions in normalised TEER after the first exposure that persisted across the exposure period (**Figure 2C,D**). Filtered-smoke groups showed intermediate trajectories between air and the corresponding unfiltered conditions. At day 6, mean TEER was higher after filtered than unfiltered CBD-dominant smoke (257.8 versus 39.5 Ω·cm²; p<0.05; **Figure 2A**). Mean TEER was also numerically higher after filtered THC-dominant smoke (187.8 versus 89.0 Ω·cm²), but this comparison was not statistically significant (**Figure 2B**).

**Figure 2.**
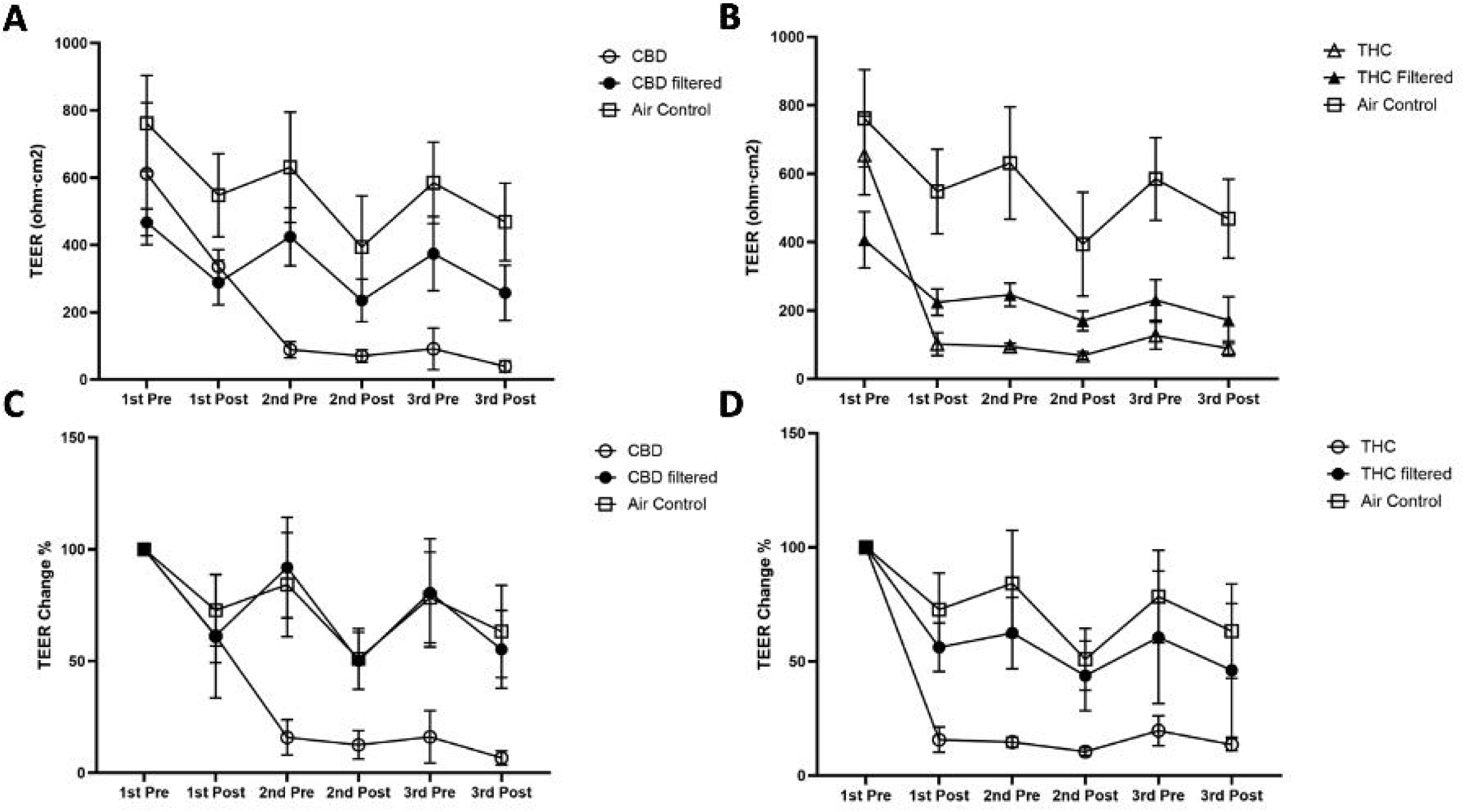
Transepithelial electrical resistance during repeated cannabis-smoke exposure. Raw TEER (A, B) and values normalised to the first pre-exposure measurement (C, D) are shown for CBD-dominant (A, C) and THC-dominant (B, D) smoke with (solid shapes) or without filtration (open shapes) with air controls. Measurements were obtained immediately before and after each exposure. Data are mean±SD for four donors (one Transwell per donor and condition); trajectories are descriptive.

At day 6, mean denuded area was 57.4% after unfiltered CBD smoke and 51.0% after unfiltered THC smoke, whereas no measurable denudation was detected in air or either filtered group (**Figure 3C**). Dark surface-associated material was visible over the filtered THC-smoke cultures at day 6 (**Figure 3A**). Apical PBS washing removed this material and revealed continuous epithelial coverage without measurable denudation. Denuded area was greater after each unfiltered smoke exposure than after air and was lower with filtration within each cannabis product (all p<0.05).

**Figure 3.**
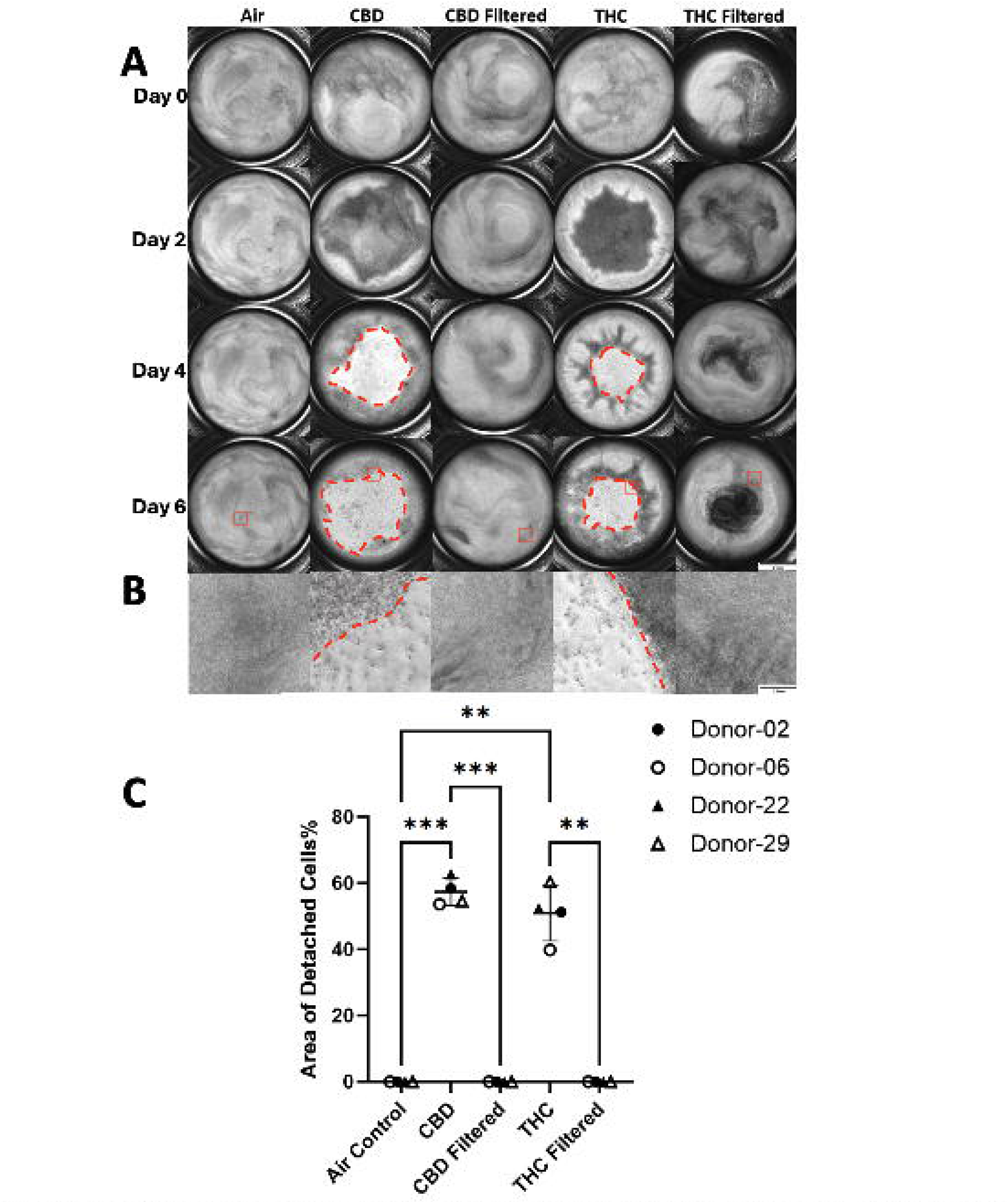
Airway epithelial cell culture surface area after repeated cannabis-smoke exposure. (A) Representative bright-field images at days 0, 2, 4 and 6; dashed outlines indicate denuded regions. (B) Enlarged regions from red boxes in panel A at Day-6. (C) Day-6 denuded area as a percentage of Transwell membrane area. Points represent donors (n=4; one Transwell per donor and condition); bars show mean±SD. Data were analysed by one-way repeated-measures ANOVA with Geisser-Greenhouse correction and Bonferroni-adjusted prespecified comparisons. *p<0.05. Scale bars: 2 mm (A) and 0.2 mm (B).

### Filtration attenuated cannabis-smoke-induced epithelial cell death

Ethidium homodimer-1-positive area was 29.1% in air controls, 91.2% after unfiltered CBD smoke and 74.5% after unfiltered THC smoke (**Figure 4B**). The corresponding filtered CBD and THC values were 38.5% and 36.0%. Each unfiltered smoke exposure increased ethidium-positive area relative to air, and filtration reduced this area within each cannabis product (all p<0.05). No pre-specified comparison in LDH release was statistically significant although the trends in data followed observations for ethidium homodimer-1-positive area (**Figure 4C**).

**Figure 4.**
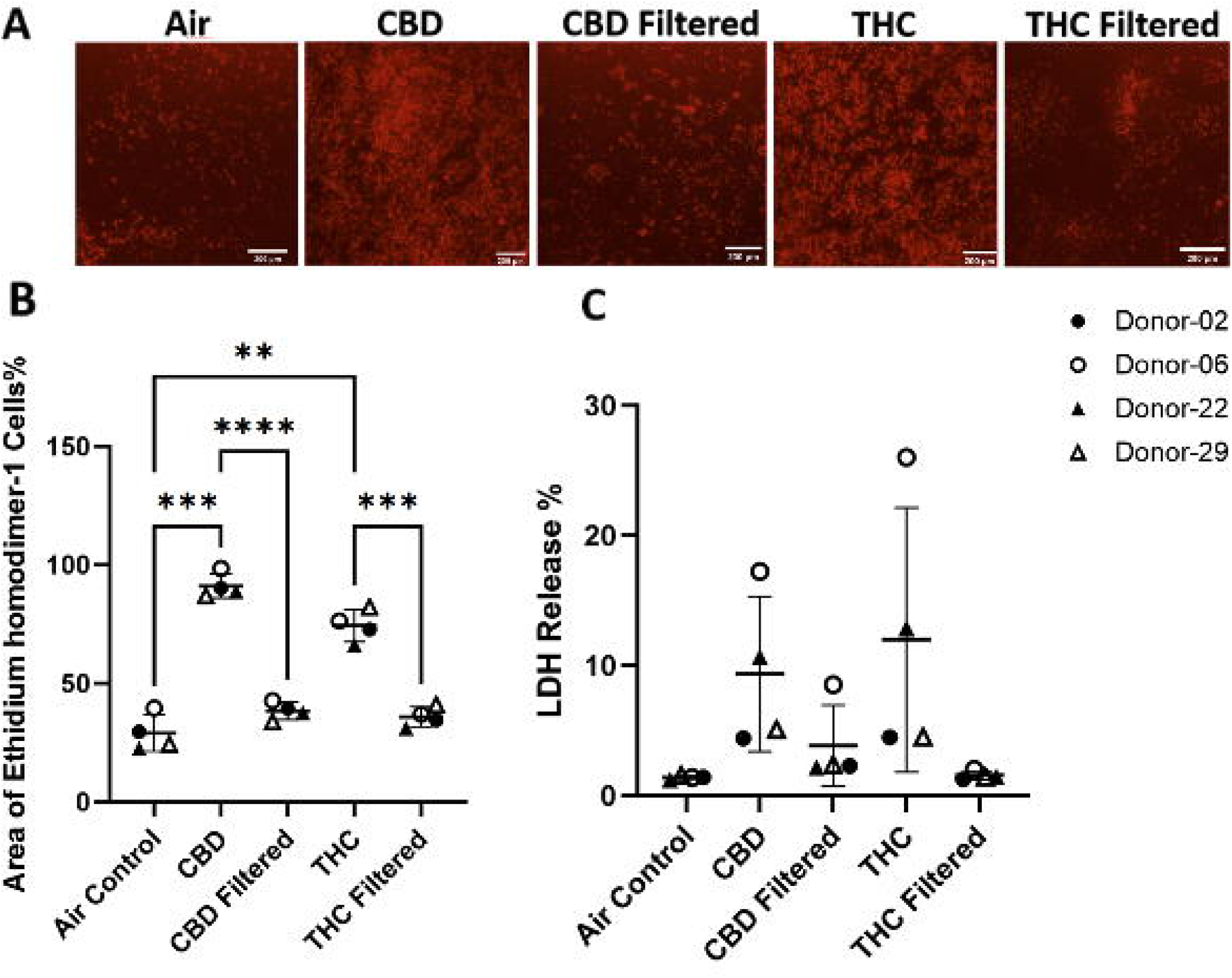
Cellular injury after repeated cannabis-smoke exposure. (A) Representative Day-6 ethidium homodimer-1 fluorescence images; scale bar, 200 µm. (B) Ethidium-positive area as a percentage of imaged membrane area (n=4; one Transwell per donor and condition). (C) Basolateral LDH release 48 h after the first exposure (n=4). Points represent donors; bars show mean±SD. Data were analysed by one-way repeated-measures ANOVA with Geisser-Greenhouse correction and Bonferroni-adjusted prespecified comparisons. *p<0.05.

### Measured soluble cannabinoid concentrations in capture medium did not differ significantly with filtration

Mean soluble CBD remained measurable in capture medium after filtered and unfiltered CBD-dominant joint combustion, with mean concentrations of 322.9 and 394.0 ng·mL□¹, respectively (filtered n=11; unfiltered n=8; **Figure 5A**). Mean soluble THC concentrations after filtered and unfiltered THC-dominant combustion were 229.9 and 194.3 ng·mL□¹, respectively (filtered n=7; unfiltered n=8; **Figure 5B**). No statistically significant differences were detected between filtered and unfiltered combustion for either analyte measured.

**Figure 5.**
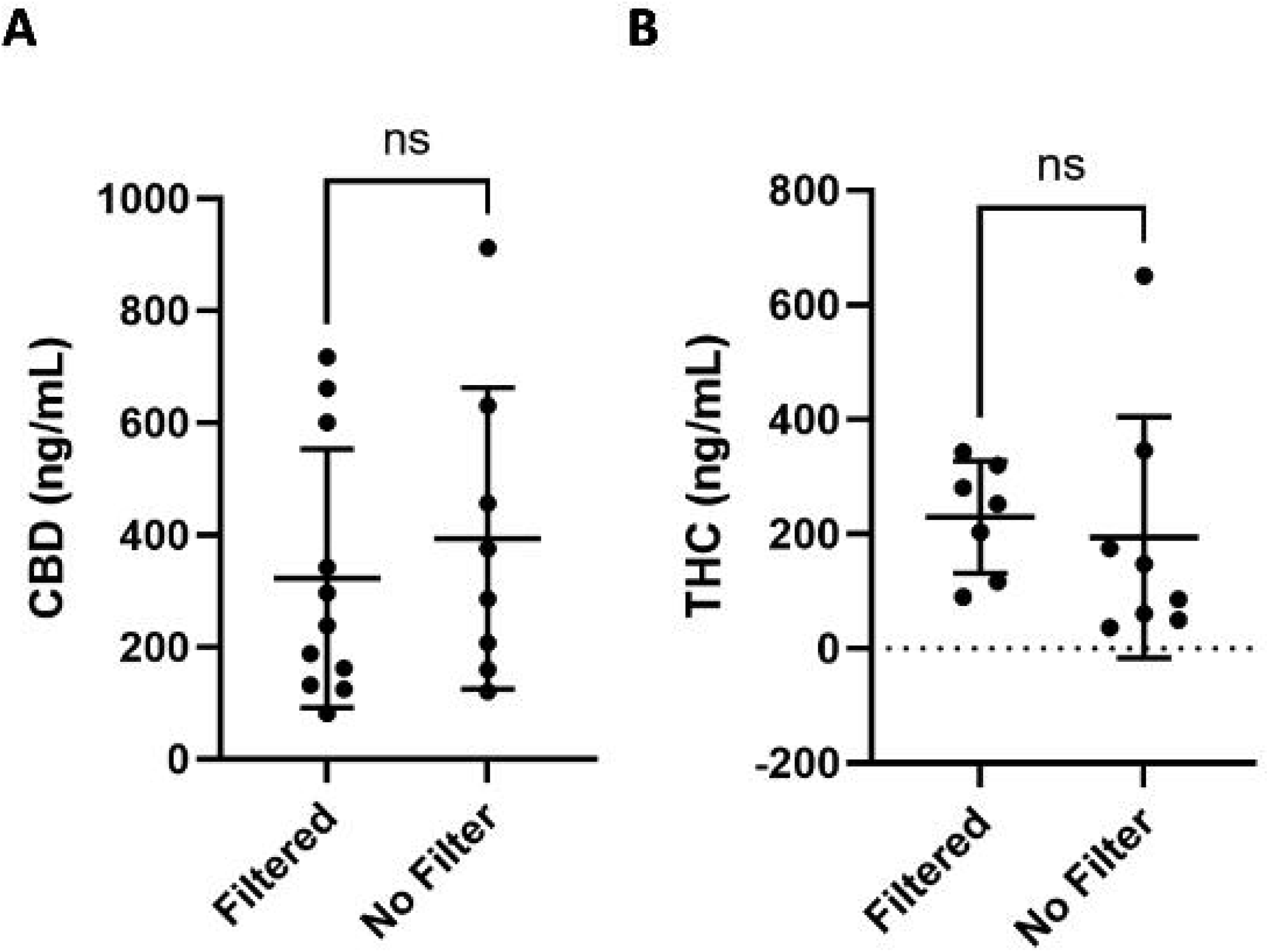
Soluble cannabinoids captured in cell culture medium after combustion of an entire cannabis joint. (A) CBD recovered following filtered (n=11) and unfiltered (n=8) CBD-dominant joint combustion. (B) Δ9-THC recovered following filtered (n=7) and unfiltered (n=8) THC-dominant joint combustion. Points represent independent 0.75 g joints entirely combusted. Data are presented as individual values with mean±SD. Filtered and unfiltered conditions were compared using two-sided unpaired Welch’s t-tests. ns, not significant.

### Filtration modified pre-specified epithelial alarmin and death-associated mediator responses

The five pre-specified mediator outcomes were IL-33, TSLP, soluble Fas, TRAIL and HMGB1 (**Figure 6A–E**). Unfiltered CBD- and THC-dominant smoke increased IL-33, soluble Fas and TRAIL relative to air (all p<0.05; **Figure 6A,C,D**). Filtration reduced soluble Fas and TRAIL after both THC and CBD-dominant products and reduced IL-33 after THC-dominant product combustion (all p<0.05). No pre-specified comparison was statistically significant for TSLP or HMGB1 (**Figure 6B,E**).

**Figure 6.**
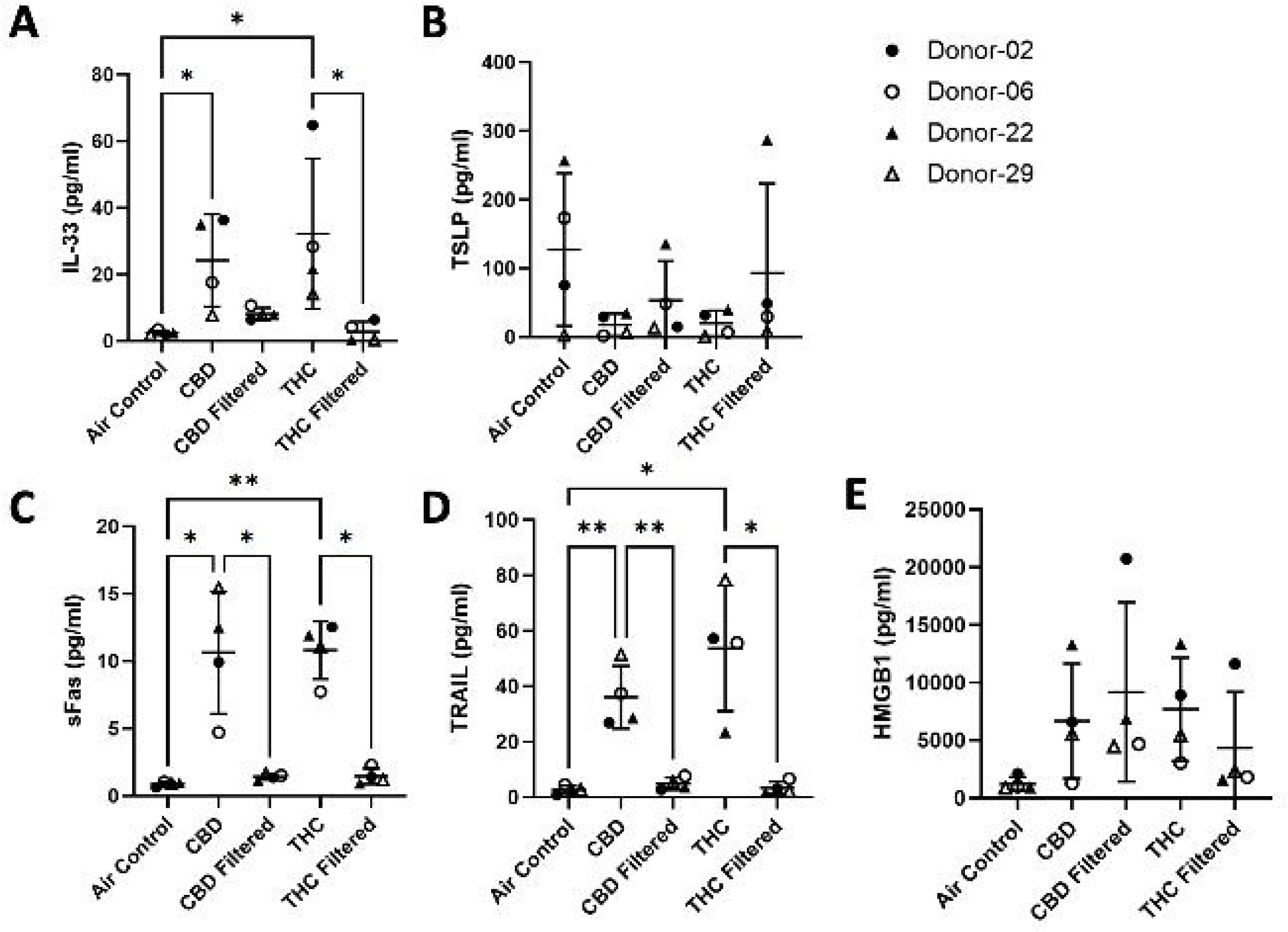
Basolateral chemokine/cytokine production 48 h after the first exposure. (A) IL-33, (B) TSLP, (C) soluble Fas (sFas), (D) TRAIL and (E) HMGB1. Points represent donors (n=4); plots show mean±SD. Statistics were performed on log normal transformed levels with one-way repeated-measures ANOVA with Geisser-Greenhouse correction and Bonferroni-adjusted pre-specified comparisons (see methods). *p<0.05.

### Filtered cannabis smoke altered epithelial marker^high^ cell fractions

Flow-cytometric analysis was restricted to air and filtered-smoke conditions because unfiltered cultures yielded insufficient cells for practical experiments with reliable live-cell analyses. Acetylated alpha-tubulin^high^ fractions decreased from 33.9% in air controls to 0.9% after filtered CBD and 0.4% after filtered THC smoke (**Figure 7B**, both p<0.05). The MUC5AC^high^ fraction decreased from 38.4% in air controls to 15.7% after filtered CBD smoke (p<0.05), and did not differ significantly after filtered THC smoke. The cytokeratin 5^high^ fraction increased from 50.4% in air controls to 68.7% after filtered CBD smoke (p<0.05), and did not differ significantly after filtered THC smoke. SCGB1A1^high^ fractions did not differ significantly between either filtered condition and air.

**Figure 7.**
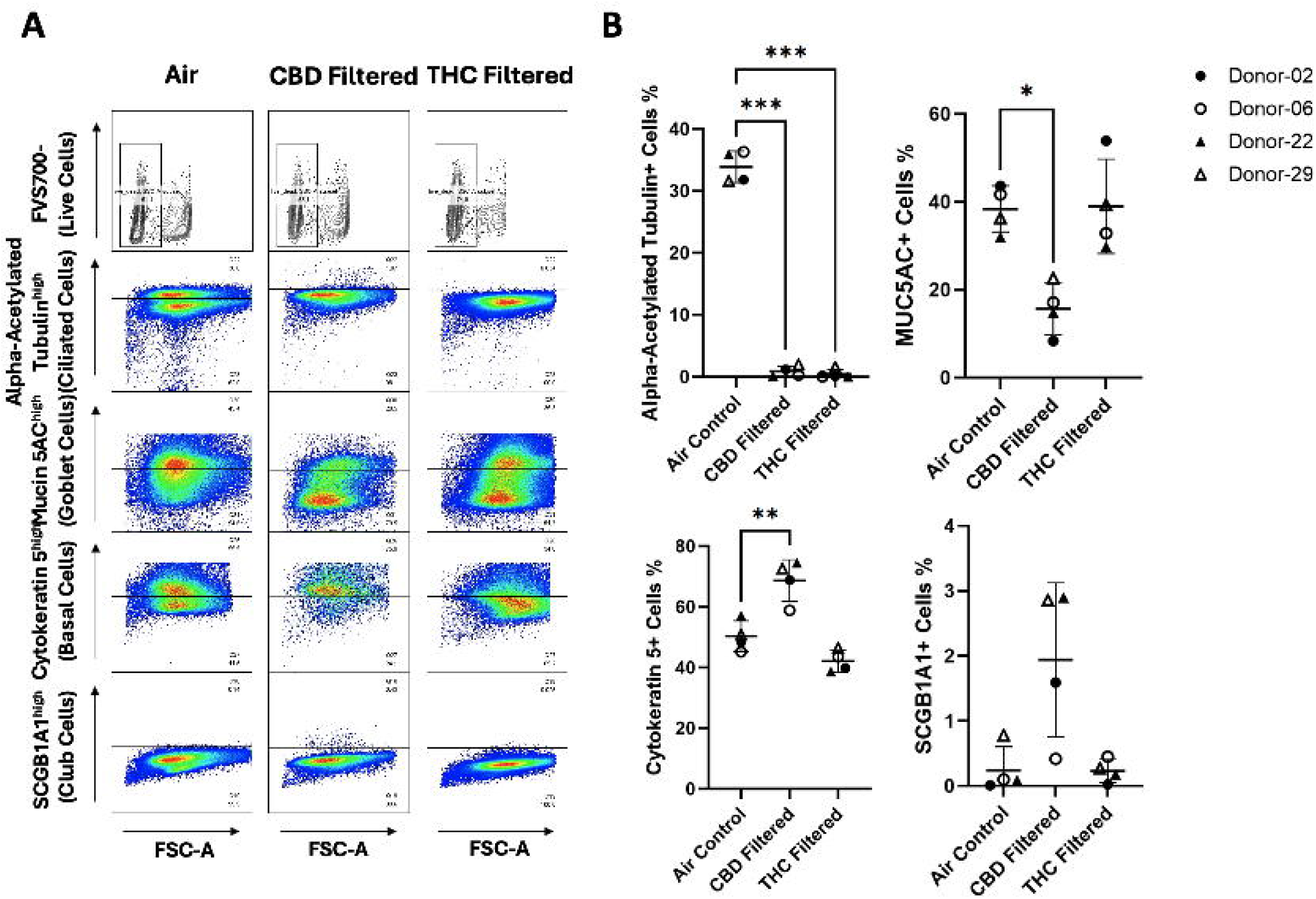
Flow-cytometric analysis of epithelial markers after repeated filtered cannabis-smoke exposure. (A) Representative sequential live-cell and singlet gating and intracellular staining for SCGB1A1, cytokeratin 5, acetylated α-tubulin and MUC5AC in air-control, filtered CBD-smoke and filtered THC-smoke conditions. (B) Acetylated α-tubulin-, MUC5AC-, cytokeratin 5- and SCGB1A1-high fractions among viable singlets were used to identify ciliated epithelial cells, goblet cells, primary basal cells and club cells, respectively. To generate sufficient cells for cytometry, cells from four Transwell inserts were pooled per donor and condition (n=4). Points represent individual donors; bars show mean±SD. Data were analysed separately for each marker by one-way repeated-measures ANOVA with Geisser-Greenhouse correction and Bonferroni-adjusted comparisons with air. *p<0.05.

## Discussion

Cellulose-acetate filters are widely used in commercial tobacco cigarettes, yet they have not been explored as a harm-reduction approach for cannabis combustion products. We evaluated cellulose-acetate filters in an *in vitro* model of primary human airway epithelial cells cultured under ALI conditions. Filtration attenuated epithelial denudation and cell-death staining after smoke exposure from both THC- and CBD-dominant cannabis. Day-6 barrier resistance was higher with filtration for CBD-dominant smoke, whereas the corresponding THC comparison was not statistically significant. These changes were observed despite measurable soluble cannabinoid concentrations in capture medium not differing significantly between filtered and unfiltered combustion conditions. Cellulose-acetate filters therefore present a promising harm-reduction strategy for cannabis flower combustion, although it should be noted that like tobacco smoke, negative impacts of cannabis combustion are likely to persist even with filtration.

Cannabis combustion represents a growing public-health concern. An estimated 256 million people used cannabis worldwide in 2024, approximately 40% more than a decade earlier [1]. Legal access has expanded from national adult-use frameworks in Uruguay and Canada to 28 jurisdictions in the United States, alongside more limited non-commercial access frameworks in Malta, Luxembourg and Germany [1]. Although globally representative data on routes of administration remain limited, smoking remained the most common route among Canadian past-year consumers in 2024 (69%) [2]. Cannabis smoking is associated with both respiratory symptoms and objective airway abnormalities. Reducing or quitting frequent cannabis smoking improves cough, sputum production and wheeze [10]. Cumulative exposure has also been associated with altered lung volumes and airway conductance [23]. Bronchoscopy studies demonstrate airway inflammation despite normal spirometry [24], and the CANUCK study linked cannabis smoking with respiratory symptoms, imaging abnormalities and airway epithelial molecular changes [11]. The continued prevalence of smoked cannabis creates a need to evaluate product-level approaches that reduce airway injury.

Preservation of epithelial coverage was the clearest protective effect of filtration. Denudation and ethidium-positive area reflect loss of epithelial continuity and accumulated membrane-compromising injury, whereas TEER integrates electrical barrier function across the remaining epithelium and LDH was measured at an earlier stage of the exposure series. These differences in endpoint and timing may explain why filtration effects were evident in day-6 imaging and barrier measures for smoke from CBD-dominant flower, but less conclusive for THC-dominant flower. Previous conditioned-medium and whole-smoke models demonstrated barrier impairment after cannabis exposure [12,18]; the present findings extend that work by showing that an upstream physical modification of the joint with a cellulose acetate filter can attenuate repeated-smoke injury. Taken together, filtration protected against accumulated structural damage and improved day-6 barrier resistance after CBD-dominant smoke, whereas the THC TEER and early LDH responses were less conclusive.

Direct evidence comparing filtered and unfiltered cannabis smoke remains limited. Aloufi et al. generated cannabis smoke extract from joints containing a cellulose filter and found inflammatory responses in primary human lung fibroblasts, but the absence of an unfiltered arm prevented assessment of filter efficacy [34]. In a machine-generated emissions study, Sambiagio et al. found that a combined cellulose-activated-carbon filter reduced several classes of toxicants while the reported THC delivery efficiency was similar for joints with and without filtration [35]. Although that study used a combined cellulose-activated-carbon filter rather than the cellulose-acetate filter used here, its findings support the concept that filtration can alter toxicant emissions without proportionally changing cannabinoid recovery. A practical question was whether the reduction in epithelial injury was accompanied by lower cannabinoid recovery. In our independent smoke-capture experiments, filtration did not significantly alter soluble CBD or THC concentrations recovered in DMEM. This finding should not be interpreted as evidence of equivalent cannabinoid delivery, because the assay quantified only cannabinoids recovered in the soluble fraction after passage through the system and did not measure cannabinoids retained by the filter or remaining in uncaptured particulate material. Cannabis smoke contains substantial semi-volatile material [8], and cannabinoid composition can change during joint combustion through volatilisation, condensation and re-evaporation [25], making recovery sensitive to puffing conditions, filter loading and capture efficiency. Smoke bioactivity also does not necessarily parallel particulate mass or cannabinoid concentration [26]. The reduced epithelial injury could therefore reflect lower particle exposure or changes in unmeasured toxicants, rather than preferential preservation of cannabinoid delivery.

Filtration attenuated selected epithelial alarmin and death-associated mediator responses. Lower IL-33 after filtered smoke from THC-dominant flower is compatible with reduced epithelial danger signalling; because oxidative stress can promote IL-33 release [27,28], a lower oxidant burden is one possible but untested explanation. Parallel reductions in soluble Fas and TRAIL are consistent with modified death-associated signalling but do not identify a specific cell-death pathway [29,30]. These biochemical changes did not translate into preservation of epithelial differentiation: filtered smoke still altered epithelial marker^high^ fractions associated with ciliated, goblet and basal cell phenotypes. Repeated tobacco-smoke exposure can disrupt epithelial differentiation [15], while airway samples from habitual cannabis smokers demonstrate ciliary abnormalities, basal-cell changes and altered MUC5AC expression [11,31]. These observations indicate that reducing overt structural injury and selected injury-associated signals does not necessarily preserve the differentiated epithelial phenotype.

Flow cytometry also suggested product-specific phenotypic patterns. In comparisons with air, both filtered products markedly reduced acetylated α-tubulin^high^ fractions, while filtered CBD smoke was additionally associated with a reduced MUC5AC^high^ fraction and an increased cytokeratin 5^high^ fraction. Previous ALI experiments showed cannabis-product-dependent differences in epithelial barrier injury [18], and acute mouse exposures to products with different THC:CBD ratios produced distinct pulmonary immune responses [36]. These observations support further comparison of chemically defined CBD- and THC-dominant products.

These findings support further evaluation of filtration as a cannabis-product design strategy. The tobacco experience shows that reductions in machine-measured smoke constituents do not automatically translate into lower human risk: cellulose-acetate filters were associated with misleading perceptions of safety [19,32], filter design can selectively alter smoke chemistry and biological activity [20,21], and resistance and puffing behaviour can modify the inhaled dose [33]. Filtered cannabis products therefore require controlled development and evaluation before any safety or reduced-risk claim is considered. Regulatory frameworks should require filter material, dimensions, ventilation and resistance to be specified as product-design characteristics, and standardized testing should quantify cannabinoid delivery, particulate matter, carbonyls, volatile organic compounds and polycyclic aromatic hydrocarbons under controlled and human-relevant puffing conditions.

Future cannabis-filter studies can draw on tobacco-filter evaluations that combine controlled machine smoking, phase-specific chemical analysis and biological testing [20,21,37]. These studies show that filter performance depends on the constituent assessed, filter design and puffing conditions. Applying this approach to cannabis smoke would help link filter-associated changes in smoke composition to epithelial responses.

Further work should identify which filter-induced changes in smoke composition account for the observed reduction in epithelial injury and whether this protection persists under conditions that more closely reflect human use. Candidate analytes include terpenes, carbonyls, volatile organic compounds and polycyclic aromatic hydrocarbons. Phase-resolved mass-balance studies should quantify cannabinoids and toxicants in the gas and particulate phases of filtered and unfiltered smoke, spent filters and capture medium. Comparisons across filter materials and geometries should account for pressure drop, ventilation, filter loading and realistic puff topography. Longer exposure and recovery studies using a larger and more diverse donor pool should determine the durability of the epithelial effects. Human puff-topography and exposure-biomarker studies could then assess whether filtration translates into measurable reductions in exposure. Importantly, end-user experience with the diverse filter possibilities should remain front of mind to ensure adoption of this strategy is best aligned with the end-user interests.

Strengths of this study include donor-matched comparisons in differentiated primary epithelium, repeated whole-smoke exposure and complementary structural, biochemical, mediator and phenotypic endpoints. Interpretation is constrained by the four male donors, the use of two commercial cannabis products and a repeated-acute exposure schedule. Four of seven attempted donor cultures met the prespecified ALI quality-control criteria, illustrating the practical variability that is a common, yet underreported feature of using primary human airway epithelial cell models. The donor-matched design supports the conclusion that cellulose-acetate filtration attenuated overt epithelial injury, while epithelial phenotypic alterations persisted. These findings support partial protection in this *in vitro* model; the safety of filtered cannabis smoking in humans remains untested.

## Conclusion

Cellulose-acetate filtration attenuated epithelial denudation and cell-death staining after repeated cannabis-smoke exposure in donor-derived ALI cultures and altered selected mediator responses. Measured soluble cannabinoid concentrations in capture medium did not differ significantly between filtered and unfiltered combustion. Filtered smoke nevertheless altered epithelial marker^high^ fractions; these findings support partial protection in this model, but do not establish safer smoking in humans, which requires further pre-clinical and clinical studies that extend this body of work.

## Acknowledgements

The authors acknowledge the Canada Research Chairs program for stipend support for JAH, JM, and IS and the CIHR Postdoctoral Fellowship program for stipend support for NM.

AI tools have been used to review grammar, spelling, and structure of the manuscript.

## Take Home Message

Cellulose acetate filters added to cannabis joints reduces airway epithelial cell damage in an *in vitro* smoke exposure model system.

