## Supplementary Methods for "Filtration attenuates cannabis smoke-induced impacts on primary human airway epithelial cells"

**Lactate dehydrogenase assay**

Lactate dehydrogenase (LDH) activity in basolateral cell-culture medium was measured with the CyQUANT LDH Cytotoxicity Assay Kit (Invitrogen, cat. no. C20300, Carlsbad, CA, USA). Samples and reaction mixture (50 µL each) were incubated for 30 min at room temperature protected from light, followed by 50 µL stop solution. Absorbance was measured at 490 nm with 680 nm as the reference wavelength using a SpectraMax i3x Multi-Mode Microplate Reader (Molecular Devices, Sunnyvale, CA, USA). Spontaneous- and maximum-release controls received kit water and lysis buffer, respectively. After subtraction of the 680-nm background signal, percentage LDH release was calculated as [(experimental LDH activity − spontaneous LDH activity)/(maximum LDH activity − spontaneous LDH activity)] × 100.

**Immune mediator profiling**

Undiluted basolateral cell-culture medium was analysed by Eve Technologies (Calgary, AB, Canada) using the Human Cytokine/Chemokine Panel B 48-Plex Discovery Assay (HD48B; MILLIPLEX Human Cytokine/Chemokine/Growth Factor Panel B, MilliporeSigma, cat. no. HCYTB-60K, Burlington, MA, USA). Measurements were acquired on a Luminex 200 system (Luminex Corporation/DiaSorin, Saluggia, Italy) and processed with Bio-Plex Manager software (Bio-Rad Laboratories, Hercules, CA, USA). Concentrations were derived from analyte-specific standard curves and reported in pg·mL⁻¹; assay sensitivities ranged from 0.05 to 100 pg·mL⁻¹. The 48 analytes are listed in Supplementary Table S1. Before statistical analysis, IL-33 values reported as below the assay range (OOR <) were assigned 2.48 pg·mL⁻¹, one-half of the lowest IL-33 standard concentration (4.95 pg·mL⁻¹).

**Supplementary Table S1. Analytes included in the Human Cytokine/Chemokine Panel B 48-Plex Discovery Assay.**

| 6Ckine/CCL21 | APRIL | BAFF/BLyS | BCA-1/BLC/CXCL13 |
| --- | --- | --- | --- |
| CCL28 | CTACK/CCL27 | CXCL16 | ENA-78/CXCL5 |
| Eotaxin-2/CCL24 | Eotaxin-3/CCL26 | GCP-2/CXCL6 | Granzyme A |
| Granzyme B | HMGB1/HMG1 | I-309/CCL1 | I-TAC/CXCL11 |
| IFN-β | IFN-ω | IL-11 | IL-16 |
| IL-20 | IL-21 | IL-23 | IL-24 |
| IL-28A/IFN-λ2 | IL-29/IFN-λ1 | IL-31 | IL-33 |
| IL-34 | IL-35 | LIF | Lymphotactin/XCL1 |
| MCP-2/CCL8 | MCP-4/CCL13 | MIP-1γ/CCL15 | MIP-3α/CCL20 |
| MIP-3β/CCL19 | MPIF-1/CCL23 | Perforin | sCD137 |
| SCF | SDF-1α+β/CXCL12 | sFAS | sFASL |
| TARC/CCL17 | Thrombopoietin (TPO) | TRAIL | TSLP |

**Flow cytometry staining**

Following cell recovery and pooling, fluorescence-activated cell-sorting (FACS) buffer was prepared as DPBS containing calcium and magnesium (Gibco, cat. no. 14040133) supplemented with 2% heat-inactivated fetal bovine serum (Wisent, cat. no. 080-450). BD GolgiStop (BD Biosciences, cat. no. 554724, Franklin Lakes, NJ, USA) was added to DPBS and FACS buffer at 10 µL per 15 mL. Cells were washed in 2 mL DPBS containing GolgiStop and centrifuged at 500×g for 10 min at 4°C. The pellet was resuspended in 1 mL Fixable Viability Stain 700 (1:4000 in DPBS containing GolgiStop) and incubated for 20 min at 4°C protected from light. After washing, cells underwent a 15-min Fc-blocking step in 50 µL solution containing 10 µL BD FcR Block (BD Biosciences, cat. no. 564220) at 4°C, followed by fixation and permeabilisation in 250 µL 1× BD Cytofix/Cytoperm solution (BD Biosciences, cat. no. 554714) for 30 min at 4°C protected from light.

Fixed cells were washed in 1× Perm/Wash buffer at 300×g for 10 min at 4°C and underwent a second 15-min Fc-blocking step in Perm/Wash buffer. Intracellular antibodies (5 µL per sample) were incubated for 30 min at room temperature protected from light. Cells were then washed, resuspended in 400 µL FACS buffer without GolgiStop and stored at 4°C protected from light until acquisition within 5 days. Unstained, antibody-matched isotype and single-stained unmixing controls were processed in parallel. Matched isotype controls were used at 5 µL per sample. Single-stained antibody controls used Total Antibody Compensation Beads (Invitrogen, cat. no. A10497, Carlsbad, CA, USA). Final-panel reagents, antibodies and isotype controls are listed in Supplementary Table S2.

**Supplementary Table S2. Reagents and antibodies used for flow-cytometry staining.**

| **Reagent or marker** | **Fluorochrome/form** | **Manufacturer** | **Catalogue no.** | **Amount or dilution** |
| --- | --- | --- | --- | --- |
| Fixable Viability Stain 700 | FVS700 | BD Biosciences | 564997 | 1:4000 |
| SCGB1A1 | FITC | Santa Cruz Biotechnology | sc-365992 FITC | 5 µL/sample |
| SCGB1A1 isotype control (normal mouse IgG1) | FITC | Santa Cruz Biotechnology | sc-2855 | 5 µL/sample |
| Acetylated α-tubulin | PE | Cell Signaling Technology | 18276S | 5 µL/sample |
| Acetylated α-tubulin isotype control (rabbit IgG) | PE | Cell Signaling Technology | 5742S | 5 µL/sample |
| Cytokeratin 5 | APC | Abcam | ab224984 | 5 µL/sample |
| Cytokeratin 5 isotype control (rabbit IgG) | APC | R&D Systems (Bio-Techne) | IC1051A | 5 µL/sample |
| MUC5AC | Alexa Fluor 647 | Novus Biologicals | NBP2-32732AF647 | 5 µL/sample |
| MUC5AC isotype control (mouse IgG1) | Alexa Fluor 647 | R&D Systems (Bio-Techne) | IC002R | 5 µL/sample |
| FcR Block | Blocking reagent | BD Biosciences | 564220 | 10 µL/block |
| GolgiStop | Protein transport inhibitor | BD Biosciences | 554724 | 10 µL/15 mL buffer |
| Cytofix/Cytoperm | Fixation/permeabilisation | BD Biosciences | 554714 | 250 µL/sample |
| Total Antibody Compensation Beads | Unmixing control beads | Invitrogen | A10497 | 1 drop/control |

Flow cytometry was performed on a Cytek Aurora spectral flow cytometer (Cytek Biosciences, Fremont, CA, USA), with 200,000–600,000 ungated events acquired per sample. Data were analysed in FlowJo v10.8 (BD Biosciences, Ashland, OR, USA). Events were gated sequentially on the main cell population, SSC singlets, FSC singlets and FVS700-negative live cells (Supplementary Figure S1). Marker gates were informed by antibody-matched isotype controls and the observed donor- and condition-dependent background fluorescence (Supplementary Figure S2). Gates were set at or above the isotype-derived 98th-percentile threshold. Where a clearly bimodal distribution was present, the interpeak valley was used and the same marker gate was applied across conditions within the donor. Events above the final gate were operationally classified as SCGB1A1^high^, acetylated α-tubulin^high^, cytokeratin 5^high^ or MUC5AC^high^.

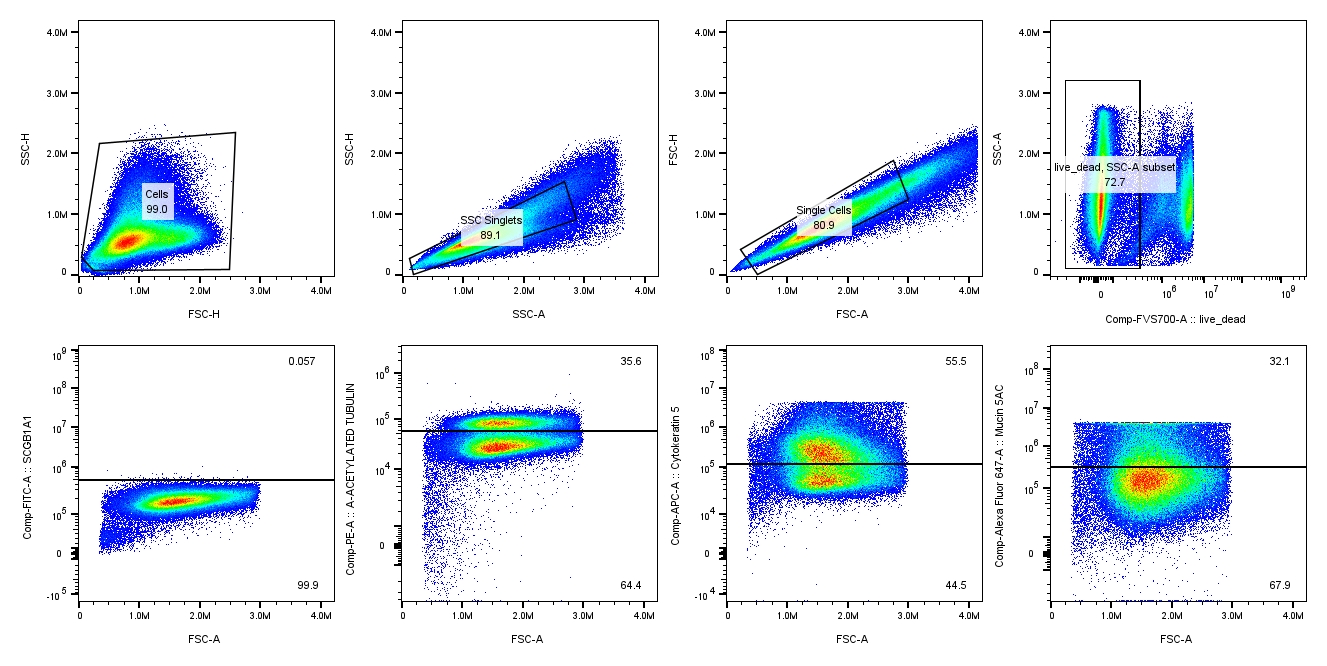

**Supplementary Figure S1.** Representative sequential gating strategy for flow-cytometric analysis. Donor-29 air-control events were gated on the main cell population, SSC singlets, FSC singlets and FVS700-negative live cells. SCGB1A1^high^, acetylated α-tubulin^high^, cytokeratin 5^high^ and MUC5AC^high^ fractions were quantified from live singlets. Numbers indicate percentages of the parent populations.

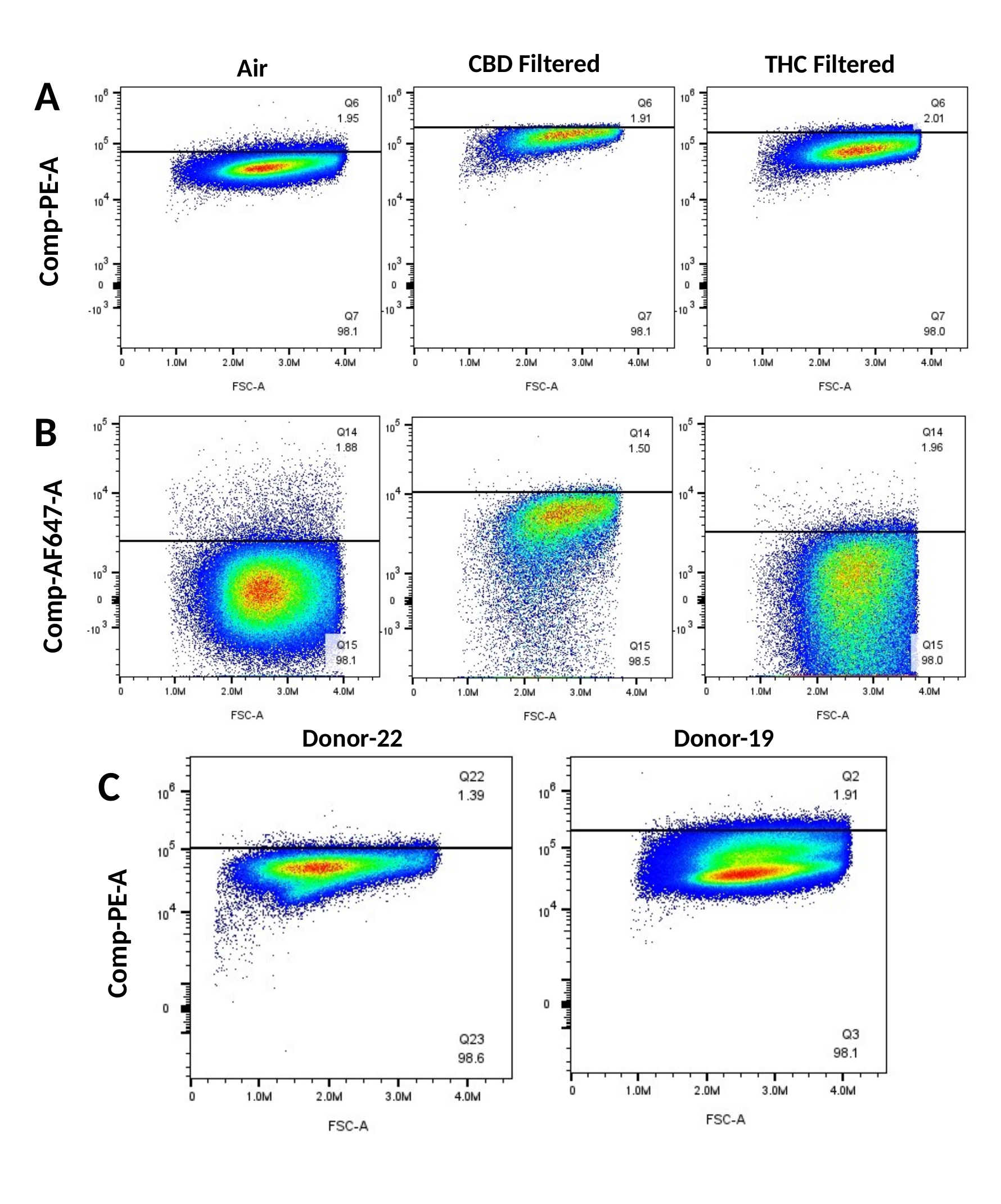

**Supplementary Figure S2.** Representative isotype-control fluorescence distributions. (A, B) PE- and AF647-conjugated isotype controls, respectively, from Donor-19 under air, filtered CBD and filtered THC conditions. (C) PE-conjugated isotype-control distributions representing the three exposure conditions in Donor-22 and Donor-19, illustrating between-donor variation in background fluorescence. Gates were positioned so that at least 98% of isotype-control events fell below the threshold. Donor-19 was used for control optimisation and was not included in the outcome analyses.

**Cannabis Smoke Profiling for Cannabinoids**

Analytical standards for CBD, Δ9-THC and 11-hydroxy-Δ9-THC and their matched d3-labelled internal standards were obtained from Sigma-Aldrich (St Louis, MO, USA). LC-MS-grade water and acetonitrile were obtained from Caledon Laboratories (Georgetown, ON, Canada), and LC-MS-grade formic acid and isopropanol from Fisher Scientific (Fair Lawn, NJ, USA). Calibrators were prepared at 1, 2, 5, 10, 25, 50, 100 and 200 ng·mL⁻¹ in 1:1 water:acetonitrile containing 20 ng·mL⁻¹ of the internal-standard mixture. Matrix quality-control samples were prepared at 100, 200 and 1000 ng·mL⁻¹ before extraction and underwent the same 10-fold dilution as study samples, producing final extract concentrations of 10, 20 and 100 ng·mL⁻¹.

Capture-medium samples were thawed and vortexed for 10 s. A 100-µL aliquot was combined with 400 µL cold acetonitrile containing 1% formic acid and the internal-standard mixture at 50 ng·mL⁻¹, vortexed for 1 min and centrifuged at 18,000×g for 15 min. A 100-µL aliquot of clarified supernatant was transferred to an HPLC vial with a glass insert, combined with 100 µL LC-MS-grade water and vortexed for 10 s before analysis. A 5-µL volume was injected into an Agilent 1290 Infinity II HPLC coupled to an Agilent 6495C iFunnel triple-quadrupole mass spectrometer. Separation used an Agilent RRHD Eclipse Plus C18 column (100×2.1 mm, 1.8 µm) maintained at 40°C; the autosampler was maintained at 10°C. Mobile phase A was 0.1% formic acid in water and mobile phase B was 0.1% formic acid in acetonitrile. The flow rate was 0.3 mL·min⁻¹ with a linear gradient from 50% to 90% B over 6 min.

Detection used positive-ion electrospray ionisation and multiple-reaction monitoring. Source settings were: drying-gas flow, 11 L·min⁻¹; drying-gas temperature, 290°C; nebuliser pressure, 25 psig; sheath-gas flow, 11 L·min⁻¹; sheath-gas temperature, 300°C; capillary voltage, 3000 V; nozzle voltage, 2000 V; and acquisition rate, 1.24 cycles·s⁻¹. Quantification was based on analyte-to-internal-standard peak-area ratios using unweighted linear regression. Calibration standards were injected at the beginning, middle and end of each analytical run, and the mean response of bracketing standards was used to quantify the corresponding samples. Calibration curves required R²>0.99 and back-calculated calibrator concentrations within ±30% of nominal; acceptable quality-control recovery was 80–120% with SD<20%. Study and quality-control samples were injected in duplicate, with a required percentage difference ≤10%, and the mean concentration was reported. Following analytical review and before statistical comparison, complete capture samples were excluded if a CBD-dominant sample contained quantifiable Δ9-THC (≥29 ng·mL⁻¹) or a THC-dominant sample contained 11-hydroxy-Δ9-THC above its method detection limit (11 ng·mL⁻¹; transition m/z 331.2 → 313.2; typical retention time 4.307 min). No analyte-specific exclusions were made. Final analyses included 11 filtered and eight unfiltered CBD-dominant samples and seven filtered and eight unfiltered THC-dominant samples. Analyte transitions, retention times, detection limits, quantification limits and quality-control recoveries are provided in Supplementary Table S3; concentrations used for Figure 5 are provided in Supplementary Data S1 (Data used for Figure 5).

**Supplementary Table S3. LC-MS/MS parameters and quality-control performance for CBD, THC and 11-hydroxy-Δ9-THC.**

| **Parameter** | **CBD** | **Δ9-THC** | **11-OH-THC** |
| --- | --- | --- | --- |
| Analyte transition (m/z) | 315.2 → 193.0 | 315.2 → 193.1 | 331.2 → 313.2 |
| d3 internal-standard transition (m/z) | 318.3 → 196.0 | 318.3 → 196.1 | 334.3 → 316.2 |
| Typical retention time (min) | 5.081 | 6.301 | 4.307 |
| Method detection limit (ng·mL⁻¹) | 7.5 | 9.7 | 11 |
| Limit of quantification (ng·mL⁻¹) | 23 | 29 | 32 |
| Low QC recovery, mean±SD (%) | 87±2 | 86±2 | 99±2 |
| Mid QC recovery, mean±SD (%) | 91±5 | 91±6 | 106±5 |
| High QC recovery, mean±SD (%) | 86±3 | 90±3 | 102±3 |
